# Intracellular reactive oxygen species (intraROS)-aided localized cell death contributing to immune responses against wheat powdery mildew pathogen

**DOI:** 10.1101/2022.10.06.511165

**Authors:** Yinghui Li, Rajib Roychowdhury, Liubov Govta, Samidha Jaiwar, Zhen-Zhen Wei, Imad Shams, Tzion Fahima

## Abstract

Reactive oxygen species (ROS) and hypersensitive response (HR) mediated cell death have long been known to play critical roles in plant immunity to pathogens. Wheat powdery mildew caused by *Blumeria graminis* f. sp. *tritici* (*Bgt*) is a destructive wheat pathogen. Here, we report a quantitative analysis of the proportion of infected cells with local apoplastic ROS (apoROS) versus intracellular ROS (intraROS) accumulation in various wheat accessions that carry different disease resistance genes (R genes), at a series of time points post-infection. The proportion of apoROS accumulation was 70-80% of the infected wheat cells detected in both compatible and incompatible host-pathogen interactions. However, intensive intraROS accumulation followed by localized cell death responses were detected in 11-15% of the infected wheat cells, mainly in wheat lines that carried nucleotide-binding leucine-rich repeat (NLR) R genes (e.g. *Pm3F, Pm41, TdPm60, MIIW72, Pm69*). The lines that carry unconventional R genes, *Pm24* (*Wheat Tandem Kinase 3*) and *pm42* (a recessive R gene), showed very less intraROS responses, while 11% of *Pm24* line infected epidermis cells still showed HR cell death, suggesting that different resistance pathways are activated there. Here, we also demonstrated that ROS could not act as a strong systemic signal for inducing high resistance to *Bgt* in wheat, although it induced the expression of pathogenesis-related (*PR)* genes. These results provide new insights on the contribution of intraROS and localized cell death to immune responses against wheat powdery mildew.

## INTRODUCTION

During host-pathogen co-evolution, plants developed multifaceted innate immunity composed of two interconnected layers of immune systems: the pathogen-associated molecular pattern (PAMP)-triggered immunity (PTI) and effector-triggered immunity (ETI) (Jones and Dangl 2006). The PTI is activated by the recognition of plant cell surface pattern recognition receptors (PRRs), conferring plants’ basic resistance to pathogens (Boutrot and Zipfel 2017; Lorang 2019). ETI responses are generally intracellular and triggered by the specific interaction between nucleotide-binding domain leucine-rich repeat (NBS-LRR)-containing receptors (NLRs) and pathogen effectors (Macho and Zipfel 2014; Zhang and Coaker 2017). Although PTI and ETI responses are triggered by different pathogen-derived molecules and crosstalk in several downstream signals, inducing defense mechanisms through reactive oxygen species (ROS), hypersensitive response (HR), plant hormones and pathogenesis-related (PR) proteins (Ali et al. 2018; Ngou et al. 2022; Yuan et al. 2021). Traditionally, particular ROS, like superoxide radicals (O_2-_), hydrogen peroxide (H_2_O_2_), and hydroxyl radicals (OH^-^) are considered as inevitably harmful by-products created during aerobic metabolism (Foyer and Noctor 2005). Moreover, ROS accumulation is also known as one of the important cellular signaling molecules playing different roles in multiple plant processes, including responses to biotic and abiotic stimuli, and plant growth development (Hasanuzzaman et al. 2020).

Respiratory burst oxidase homolog (RBOH) proteins, which are known to generate ROS in plant cells, are located on the plasma membrane (Hasanuzzaman et al. 2020; Torres et al. 2017). For example, during PTI, the recognition of flg22 or chitin via PRRs leads to transient but robust production of ROS around the apoplast (apoROS hereafter) by the membrane-bound RBOHD (Lammertz et al. 2019; Su et al. 2021; Tjamos et al. 2022). The apoROS burst around the plant cell membrane can act as an antimicrobial molecule and strengthen the plant cell wall through oxidative crosslinking (Dey et al. 2020; Jwa and Hwang 2017). The other major generation sites of ROS inside plant cells (intracellular ROS or intraROS hereafter) are mainly chloroplasts as a consequence of the disruption and imbalance of metabolic pathways, which play an important role in HR during the ETI that is probably triggered by the recognition of pathogen effectors by NLRs (Mittler et al. 2022; Qi et al. 2019; Xu et al. 2019). However, some pathogen effectors can suppress ROS accumulation involved in plant immunity (Liu et al. 2021; Ramachandran et al. 2017; Shidore et al. 2017). The production of ROS by the NADPH oxidase RBOHD is a critical early signaling event connecting PRR- and NLR-mediated immunity (Yuan et al. 2021).

HR can be morphologically defined as a type of programmed cell death (PCD), probably triggered upon pathogen recognition by NLRs, and acts as a powerful response against pathogens. The triggering of HR requires the integration of multiple signals employed by a complex regulatory mechanism. Recent studies demonstrated that the co-activation of PTI and ETI could improve NLR-mediated hypersensitive cell death response (Ngou et al. 2021). After the generation of ROS, it contributes to HR by activating a signaling cascade for PCD at the infection site, while inducing the expression of *PR* genes (Jones and Dangl 2006; Soliman et al. 2021). PCD was previously thought to be the outcome of ROS directly killing cells via oxidative stress, which is now considered to be a result of ROS triggering complex physiological or programmed pathways (Király et al. 2021; Mittler 2017). A recent study showed that ROS homeostasis mediated by MPK4 (protein kinase) and SUMM2 (NB-LRR protein) determines synergid cell death (Völz et al. 2022), suggesting that ROS plays an important role in cell death.

Powdery mildew (Pm) disease is caused by the biotrophic fungal pathogen *Blumeria graminis* (DC.) E.O. Speer. f. sp. *tritici* Em. Marchal (*Bgt*), resulting in serious yield losses of wheat (*Triticum* spp.) worldwide (Savary et al. 2019). Wild emmer wheat (WEW) (*Triticum turgidum* var. *dicoccoides*), the tetraploid progenitor of cultivated bread and durum wheat, was shown to harbor novel disease resistance genes (*R*-genes) that are effective against powdery mildew (Ben-David et al. 2016; Huang et al., 2016; Nevo 2002). It is important to understand how those *R-*genes produce ROS and HR during the PTI and ETI and of interest in studies of plant immunity. Here, we conducted a quantitative analysis of apoROS, intraROS and cell death responses during the *Bgt* life cycles in various wheat accessions that carry different powdery mildew (Pm) resistance genes. The obtained results are laying the foundation for exploring the molecular signaling cascade leading to ROS-mediated cell death and associated host resistance.

## MATERIALS AND METHODS

### Plant materials and growth condition

The following wheat accessions were used in the current study. WEW accessions: G305-3M, G18-16, TD116494 (IW172), TD010009 (IW2) contain powdery mildew resistance genes *PmG3M* (*Pm69*), *PmG16* (*TdPm60*), *MlIW172*, and *Pm41*, respectively (Ben-David et al. 2010; Li et al. 2021; Wu et al. 2022; Li et al. 2020); G303-1M contains *pm42* (Hua et al. 2009); TD104088 contains a *TdPm60* and unknown *Pm* genes (Li et al. 2021); Zavitan is susceptible to *Bgt* and was used as a control. Durum wheat cultivars: Langdon, Kronos and Svevo. Bread wheat varieties: Chinese spring, Morocco and Ruta. Differential lines harboring known *Pm* resistance genes (*Pm3F, Pm13, Pm17, Pm23, Pm29, Pm32*, and *Pm24*) (Ben-David et al. 2010; Li et al. 2021). The introgression lines: LDN/G18-16/4*Ruta and LDN/G305-3M//Svevo/4*Ruta. Seeds of those accessions were obtained from the Institute of Evolution Wild Cereals Gene Bank (ICGB), at the University of Haifa, Israel (Table 1). Plants were potted and maintained in a versatile environmental test chamber with 75% humidity, 22/20°C day/night temperature regime, 12/12 h light/dark cycle, and light intensity of approximately 150 μmol m^−2^ s^−1^.

**TABLE 1.**
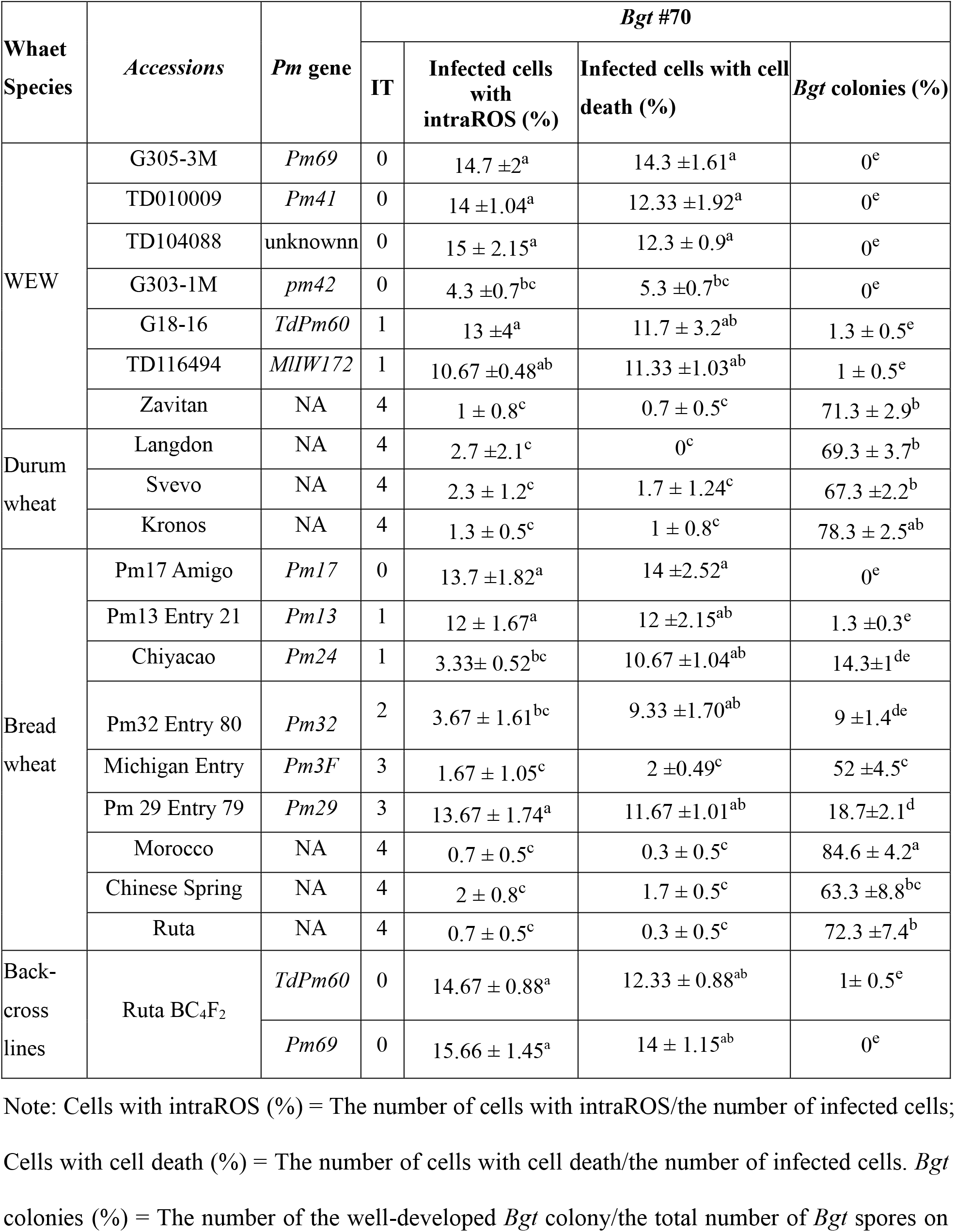
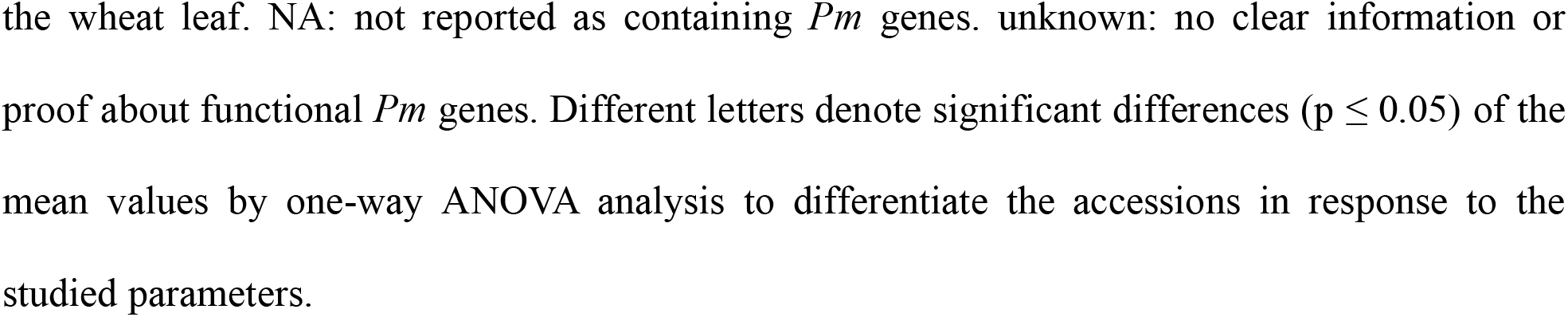
ROS and cell death response to *Bgt* #70 in different wheat species at 48 hpi.

| Wheat Species | Accessions | Pm gene | *Bgt* #70 |  |  |  |
| --- | --- | --- | --- | --- | --- | --- |
|  |  |  | IT | Infected cells with intraROS (%) | Infected cells with cell death (%) | *Bgt* colonies (%) |
| WEW | G305-3M | *Pm69* | 0 | 14.7 ± 2<sup>a</sup> | 14.3 ± 1.61<sup>a</sup> | 0<sup>e</sup> |
|  | TD010009 | *Pm41* | 0 | 14 ± 1.04<sup>a</sup> | 12.33 ± 1.92<sup>a</sup> | 0<sup>e</sup> |
|  | TD104088 | unknown | 0 | 15 ± 2.15<sup>a</sup> | 12.3 ± 0.9<sup>a</sup> | 0<sup>e</sup> |
|  | G303-1M | *pm42* | 0 | 4.3 ± 0.7<sup>bc</sup> | 5.3 ± 0.7<sup>bc</sup> | 0<sup>e</sup> |
|  | G18-16 | *TdPm60* | 1 | 13 ± 4<sup>a</sup> | 11.7 ± 3.2<sup>ab</sup> | 1.3 ± 0.5<sup>e</sup> |
|  | TD116494 | *MLIW172* | 1 | 10.67 ± 0.48<sup>ab</sup> | 11.33 ± 1.03<sup>ab</sup> | 1 ± 0.5<sup>e</sup> |
|  | Zavitan | NA | 4 | 1 ± 0.8<sup>c</sup> | 0.7 ± 0.5<sup>c</sup> | 71.3 ± 2.9<sup>b</sup> |
| Durum wheat | Langdon | NA | 4 | 2.7 ± 2.1<sup>c</sup> | 0<sup>c</sup> | 69.3 ± 3.7<sup>b</sup> |
|  | Svevo | NA | 4 | 2.3 ± 1.2<sup>c</sup> | 1.7 ± 1.24<sup>c</sup> | 67.3 ± 2.2<sup>b</sup> |
|  | Kronos | NA | 4 | 1.3 ± 0.5<sup>c</sup> | 1 ± 0.8<sup>c</sup> | 78.3 ± 2.5<sup>ab</sup> |
| Bread wheat | Pm17 Amigo | *Pm17* | 0 | 13.7 ± 1.82<sup>a</sup> | 14 ± 2.52<sup>a</sup> | 0<sup>e</sup> |
|  | Pm13 Entry 21 | *Pm13* | 1 | 12 ± 1.67<sup>a</sup> | 12 ± 2.15<sup>ab</sup> | 1.3 ± 0.3<sup>e</sup> |
|  | Chiyacao | *Pm24* | 1 | 3.33 ± 0.52<sup>bc</sup> | 10.67 ± 1.04<sup>ab</sup> | 14.3 ± 1<sup>de</sup> |
|  | Pm32 Entry 80 | *Pm32* | 2 | 3.67 ± 1.61<sup>bc</sup> | 9.33 ± 1.70<sup>ab</sup> | 9 ± 1.4<sup>de</sup> |
|  | Michigan Entry | *Pm3F* | 3 | 1.67 ± 1.05<sup>c</sup> | 2 ± 0.49<sup>c</sup> | 52 ± 4.5<sup>c</sup> |
|  | Pm 29 Entry 79 | *Pm29* | 3 | 13.67 ± 1.74<sup>a</sup> | 11.67 ± 1.01<sup>ab</sup> | 18.7 ± 2.1<sup>d</sup> |
|  | Morocco | NA | 4 | 0.7 ± 0.5<sup>c</sup> | 0.3 ± 0.5<sup>c</sup> | 84.6 ± 4.2<sup>a</sup> |
|  | Chinese Spring | NA | 4 | 2 ± 0.8<sup>c</sup> | 1.7 ± 0.5<sup>c</sup> | 63.3 ± 8.8<sup>bc</sup> |
|  | Ruta | NA | 4 | 0.7 ± 0.5<sup>c</sup> | 0.3 ± 0.5<sup>c</sup> | 72.3 ± 7.4<sup>b</sup> |
| Back-cross lines | Ruta BC<sub>4</sub>F<sub>2</sub> | *TdPm60* | 0 | 14.67 ± 0.88<sup>a</sup> | 12.33 ± 0.88<sup>ab</sup> | 1 ± 0.5<sup>e</sup> |
|  |  | *Pm69* | 0 | 15.66 ± 1.45<sup>a</sup> | 14 ± 1.15<sup>ab</sup> | 0<sup>e</sup> |

Note: Cells with intraROS (%) = The number of cells with intraROS/the number of infected cells; Cells with cell death (%) = The number of cells with cell death/the number of infected cells. *Bgt* colonies (%) = The number of the well-developed *Bgt* colony/the total number of *Bgt* spores on
the wheat leaf. NA: not reported as containing *Pm* genes. unknown: no clear information or proof about functional *Pm* genes. Different letters denote significant differences ( $p \leq 0.05$ ) of the mean values by one-way ANOVA analysis to differentiate the accessions in response to the studied parameters.

### *Bgt* inoculation and disease assessment

Two *Bgt* isolates (#70 and #SH) were regularly maintained as pure cultures in our laboratory for phenotyping tests. *Bgt* #70 was collected from *T. aestivum*; *Bgt* #SH was collected from *T. dicoccoides* (Ben-David et al. 2010; Li et al. 2021). of Pathogen inoculation, incubation conditions, and disease assessment were performed as in our previous report (Li et al. 2021). The reaction to *Bgt* inoculation was examined visually based on the disease progression on the leaf surface, and infection types (ITs) were recorded based on a scale of 0-4, where 0 represents no visible symptoms, 0; for necrotic flecks (HR), and values of 1, 2, 3 and 4 for highly resistant, resistant, susceptible and highly susceptible reactions, respectively (Xie et al. 2012).

### Evaluation of ROS accumulation and cell death

ROS accumulation and cell death were evaluated in the *Bgt* inoculated wheat leaves at different time points using previously published histochemical staining methods with minor modifications (Thordal-Christensen et al. 1997; van Wees 2008). In brief, ROS was visualized using the 3,3’-diaminobenzidine (DAB) (Sigma-Aldrich, USA) staining method in which the inoculated leaves were treated with 0.1% w/v DAB solution (pH 3.8) followed by 15 minutes of incubation in 60 kPa vacuum pressure at room temperature and 8 hours of incubation at 28° C without vacuum pressure. Then the leaf samples were decolorized in 96% (v/v) ethanol for 2 days and stained with 0.6% w/v Coomassie Brilliant Blue G250 (Sigma-Aldrich, Germany) for 2 minutes and then immediately washed with sterile water, and stored in 50% glycerol for microscopic observation. Cellular ROS and fungal structures were observed under a fluorescence microscope Leica DMi8 (Leica Microsystems, Germany) by using 2′,7′-dichlorofluorescein diacetate (H_2_DCFDA) (Sigma-Aldrich, USA) as described in Yuan et al. (2021). Cell death was assessed by 0.25% Trypan blue (Biological Industries, Israel) staining in which the treated leaves were stained for 5 minutes by incubating in boiling water, bleaching with 2.5 g mL^-1^ Chloral hydrate (Sigma-Aldrich, USA) solution for 2 days, and then stored in 50% glycerol (Sigma-Aldrich, USA) for microscopy. ROS (reddish-brown coloration) and cell death (blue coloration) were examined in the leaf tissues under a stereo microscope Zeiss Axio Imager M2 (Carl Zeiss, Germany). All microscopic experiments were repeated thrice, with at least three biological replicates each time. For the calculation of the ratios of ROS and cell death, three leaves were collected as biological replicates and observed at least 100 infected cells per leaf for identifying the ROS and cell death responses.

### Treatment with H_2_O_2_ as an external stimulus

Langdon leaves of two-week-old seedling were excised into three segments, sprayed by water (control) or 1% H_2_O_2_ solutions. Both solutions were added with 0.02% Dimethyl sulfoxide (DMSO) (Sigma-Aldrich, Germany) and 0.0001% Tween^®^ 20 (Sigma-Aldrich, USA). After spraying, the leaf segments were cultured for 4 hours on absorbent paper in square Petri dishes (12 cm X 12 cm) containing the water or 1% H_2_O_2_ solutions. Then the leaves were soaked up using absorbent paper and transferred into another square Petri dish containing 8 g L^-1^ agar with 50 mg L^-1^ benzimidazole (Sigma-Aldrich, USA) for 24 hours and finally transferred to new Petri dishes and infected with *Bgt* #70.

### RNA extraction and quantitative Reverse Transcription PCR (qRT-PCR)

Wheat leaf samples of G305-3M and Langdon were inoculated with *Bgt* #70 and sampled at different time points post inoculation (0, 3, 6, 9, 12, 16, 24, 36, 48, and 72 hpi). Non-inoculated wheat leaves were used as control. Wheat leaf samples of G305-3M and Langdon were sampled at 24 hours after treatment with 1% H_2_O_2_ solutions as described before, while H_2_O treatment was used as a control. Total RNA extraction was performed using RNeasy Plant Mini Kit (Qiagen, Germany) followed by cDNA synthesis using qScript™ cDNA Synthesis Kit (Quantabio, USA). Gene-specific primers of the *PR* genes and the housekeeping gene *Actin* are listed in Table S1. The qRT-PCR amplifications were performed with SYBR Green FastMix (Quantabio, USA) and PCR amplification was performed with StepOne thermocycler (Applied Biosystem, USA). The qRT-PCR program were as previously described (Li et al., 2021). Transcript levels are expressed as linearized fold-*Actin* levels calculated by the formula 2^(*Actin CT*-*Target* CT)^ method ± standard error of the mean (SEM). All the reactions were performed in triplicates and each reaction represents a mixed pool of three wheat leaves.

### Statistical analysis

Statistical analysis was performed using JMP^®^ version 16.0 statistical packages (SAS Institute, USA). Multiple comparisons between the genotypes for apoROS, intraROS and cell death were calculated by one-way analysis of variance (ANOVA) and Tukey-Kramer post-hoc test (for the significant ANOVA).

## RESULTS

Previously, we have identified and mapped two *Pm* resistance genes derived from WEW, namely *PmG3M* (designated hereafter as *Pm69*) (Xie et al. 2012) and *PmG16* (designated hereafter as *TdPm60*) (Li et al. 2021). The mapping populations constructed by crossing these resistant WEWs with the susceptible *T. durum* cv. Langdon, segregated for a single dominant *Pm* gene, each, showing that Langdon did not contain any functional *Pm* gene to *Bgt* isolate #70. The main motivation for the current study was to characterize ROS accumulation and HR-mediated cell death in these three lines and compare them with the response of other lines that carry different *Pm* resistance genes.

### Compatible, incompatible and partially incompatible interactions in the *Bgt*-wheat pathosystem

The macroscopic observation of fungal colonies and symptoms developed in Langdon, G305-3M and G18-16 infected with *Bgt #*70 were documented during 1-10 days post-infection (dpi) (Fig. 1). Initiation of small *Bgt* colonies was observed in Langdon already at 4 dpi, then they rapidly developed into massive fungal growth covering almost all leaf area (IT=4, highly susceptible). On the contrary, no visible disease symptoms were detected in the G305-3M (IT=0, fully resistant). In G18-16, no visible disease symptoms were detected till 6 dpi, but after that, small *Bgt* colonies could be visible and slowly developed (IT=1, partial resistance).

**Fig. 1.**
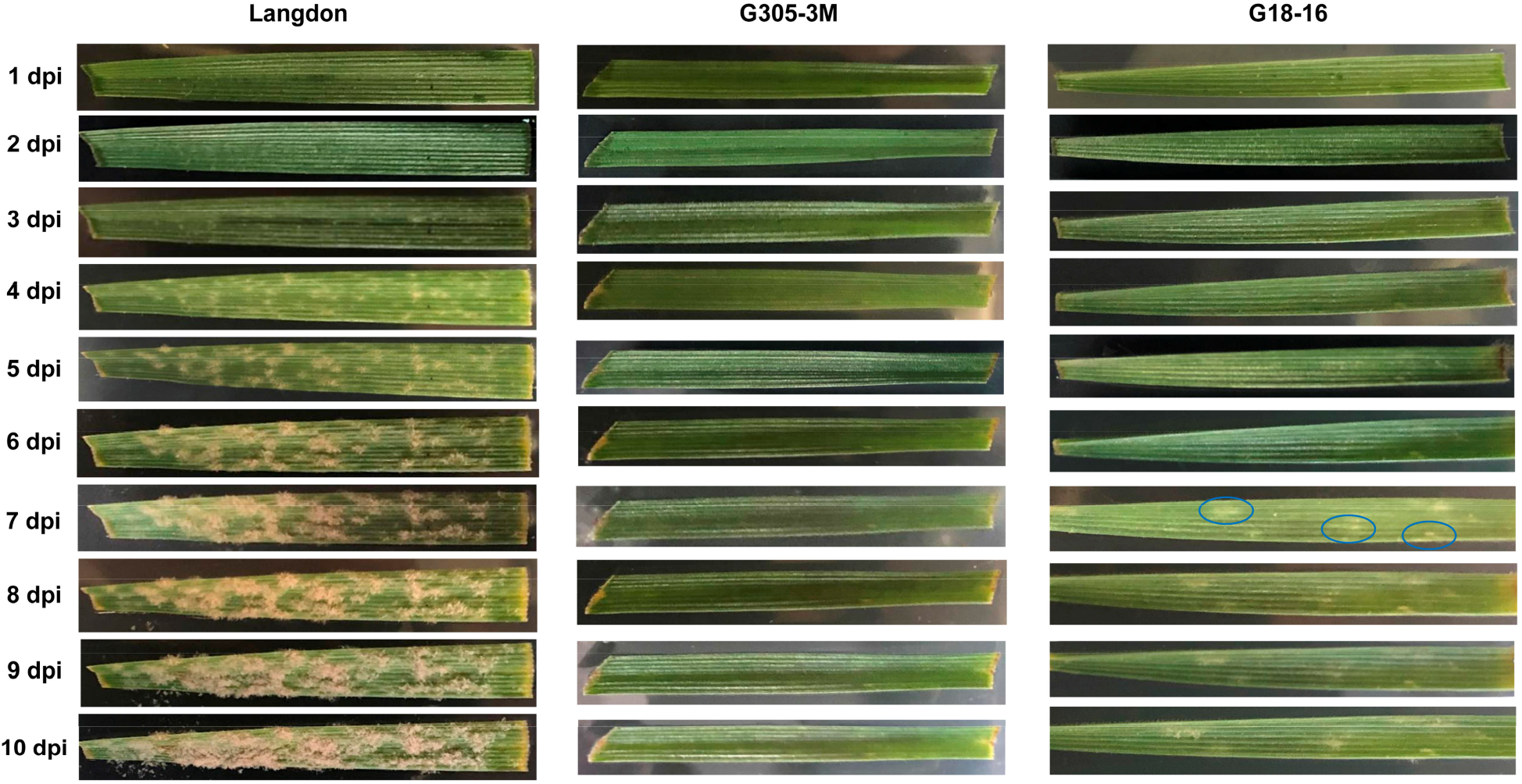
Macroscopic observation of *Bgt* #70 symptoms on young leaves of susceptible *T. durum* Langdon and two resistant WEW lines (G305-3M G18-16). The same leaves were photographed from 1 to 10 days post *Bgt* #70 infection (dpi). The blue ellipses indicate the small *Bgt* colonies on the partial resistant WEW G18-16 leaves.

### ROS accumulation is induced by powdery mildew in both compatible and incompatible *Bgt*-wheat interactions

We observed a whole asexual cycle of *Bgt* #70 developed in Langdon up to 120 hours post-infection (hpi). A germinating *Bgt* conidia (Con) formed two types of germ tubes - a primary germ tube (PGT) and a secondary germ tube (SGT) at 6 hpi, and the elongated SGT further differentiated into swollen appressoria (App) (Fig. 2). At 12 hpi, most of the App formed typical apical hooks (infection pegs) which further produced bulb-like haustorial primordia (HP) inside the host epidermal cells at 16 hpi. Subsequently, typical digitate processes (DP) of mature haustoria (MH) were visible at 24 hpi, and the formation of secondary hyphae was observed at 36 hpi. From 72 to 96 hpi, extensive hyphal growth and repeated penetration from hyphal appressoria have occurred. At 120 hpi, massive conidiophores were produced, ready to start new disease cycles (Fig. 2j).

**Fig. 2.**
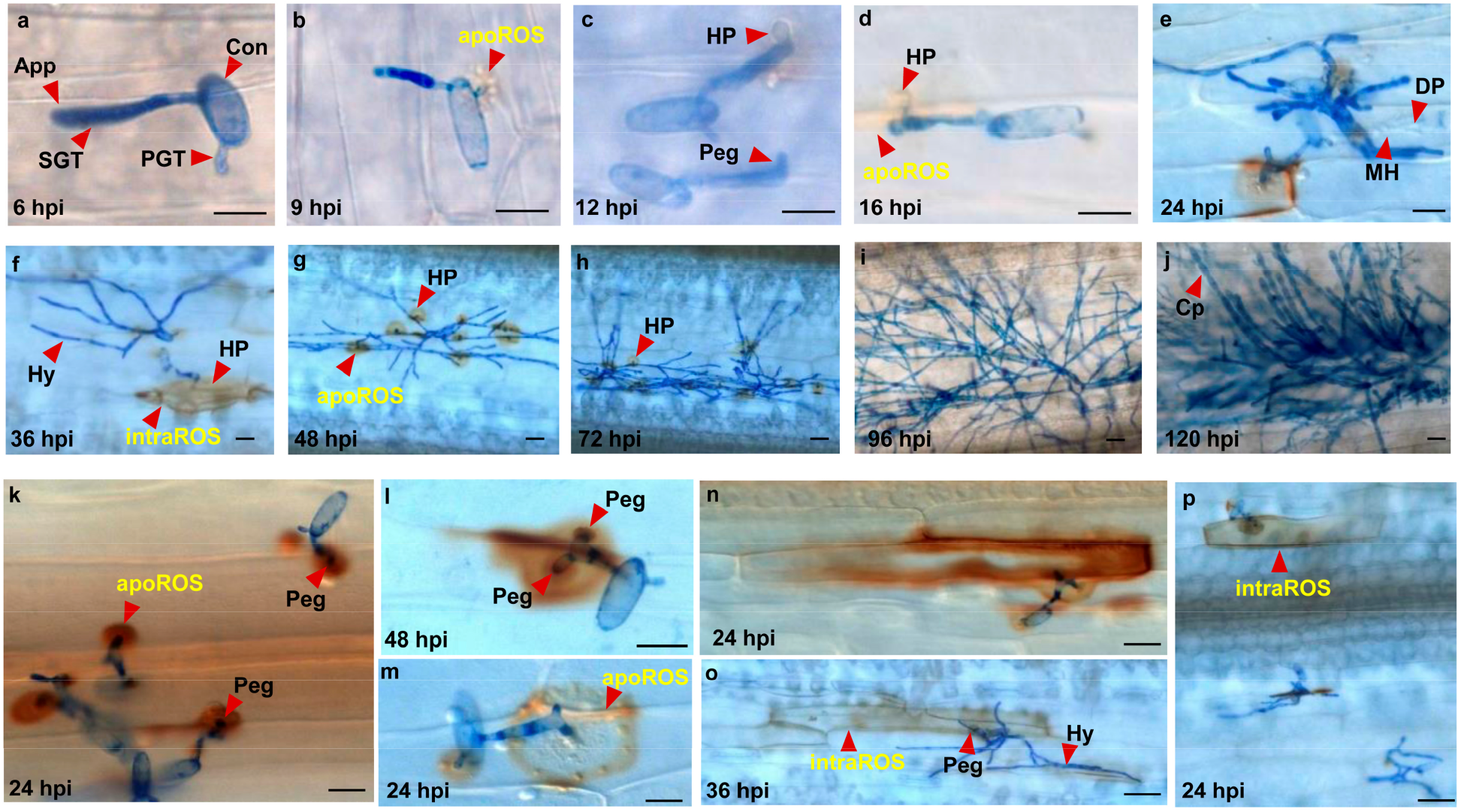
Representative micrograph of ROS accumulation during the asexual disease cycle of *Bgt* #70 in the susceptible Langdon. (**a-j**) The process of *Bgt* development and ROS accumulation from 6 to 120 hpi. (**k-p**) The different types of ROS accumulation in Langdon. App, appressorium; Con, conidium; Cp, conidiophores; DP, digitate processes (finger-like projections); MH, mature haustorium; HP, haustorial primordium; Hy, hyphae; PGT, primary germ tube; SGT, secondary germ tube. Scale bars = 20 μm.

In Langdon, most of the infected cells were found without visible ROS accumulation (Fig. S1). Nevertheless, we still found some ROS accumulation in a few epidermal cells. As shown in Fig. 2, the visible ROS formation and accumulation started at 9 hpi around the PGT, and it also could be identified around the infection peg between the time-points of 12-16 hpi (Fig. 2b-d). No ROS accumulation was observed around the MH (Fig. 2e), but the ROS could be detected around the HP during 16-72 hpi (Fig. 2d-h). Most of the ROS accumulation occurred as a halo at the penetration points of PGT and infection pegs probably located at the apoplast outside of the cell membrane (apoROS) (Fig. 2k-m). We also observed a small incidence of cells showing more extensive ROS accumulation inside the epidermal cells (intraROS) between 24-36 hpi (Fig. 2f, n, o, p).

In both G305-3M and G18-16, the *Bgt* spores germinated and developed normally from 6 to 12 hpi without any histochemical difference compared to Langdon (Fig. 3a-c, k-m). At 16 hpi, the *Bgt* gradually started to invade the epidermis host cell (Fig. 3d), however, not a single MH was detected in G305-3M, though some HP were visualized (Fig. 3e, f). In G18-16, a few *Bgt* small colonies were detected with developed hyphae (Hy) and MH during 48-120 hpi, but the disease progression was very slow (Fig. 1, 3q-t, Table 1). From 16 hpi, higher numbers of plant cells were detected with intraROS accumulation in both G305-3M and G18-16 (Fig. 3d-j, n-t), than in Langdon (Fig. 2d-j, Table 1).

**Fig. 3.**
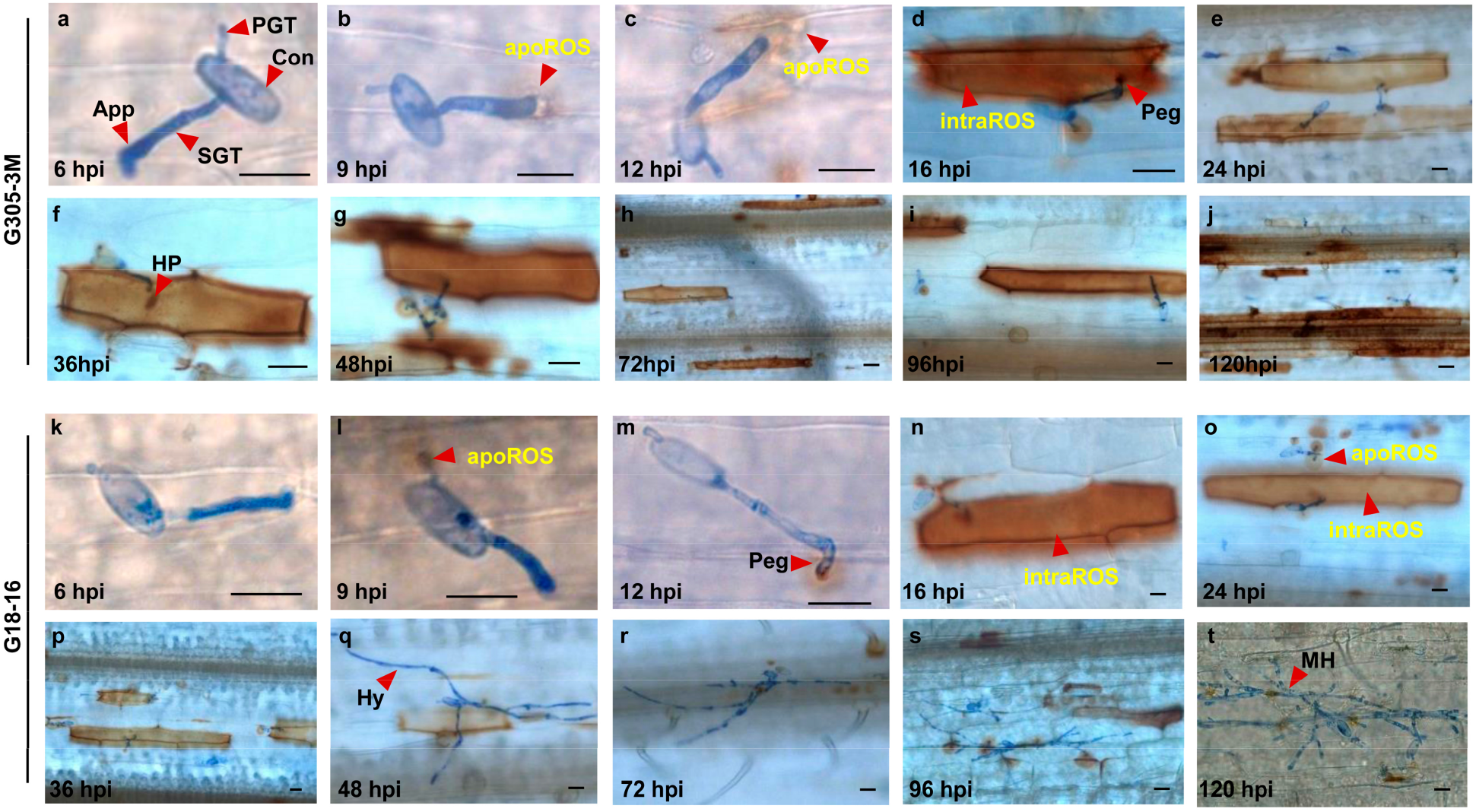
Representative micrograph of ROS accumulation during the asexual disease cycle of *Bgt* #70 in the resistant WEW lines G305-3M (**a-j**) and G18-16 (**k-t**). The process of *Bgt* development was followed from 6 to 120 hpi. App: appressorium; Con: conidium; MH, mature haustorium; PGT, primary germ tube; SGT, secondary germ tube. Scale bars = 20 μm.

### The cell death response following intraROS accumulation

In Langdon, no obvious cell death was found (Fig. 4a-e). In G305-3M and G18-16, some necrosis sites were noticed already at 16 hpi and continued to spread at 24 hpi (Fig. 4g-m and Fig. 3b, c, l, m) and 36 hpi (Fig. 4i, n), while fully developed cell death responses can be clearly seen at 48 hpi (Fig. 4j, o), and coincided with the intraROS accumulation (Fig. 3).

**Fig. 4.**
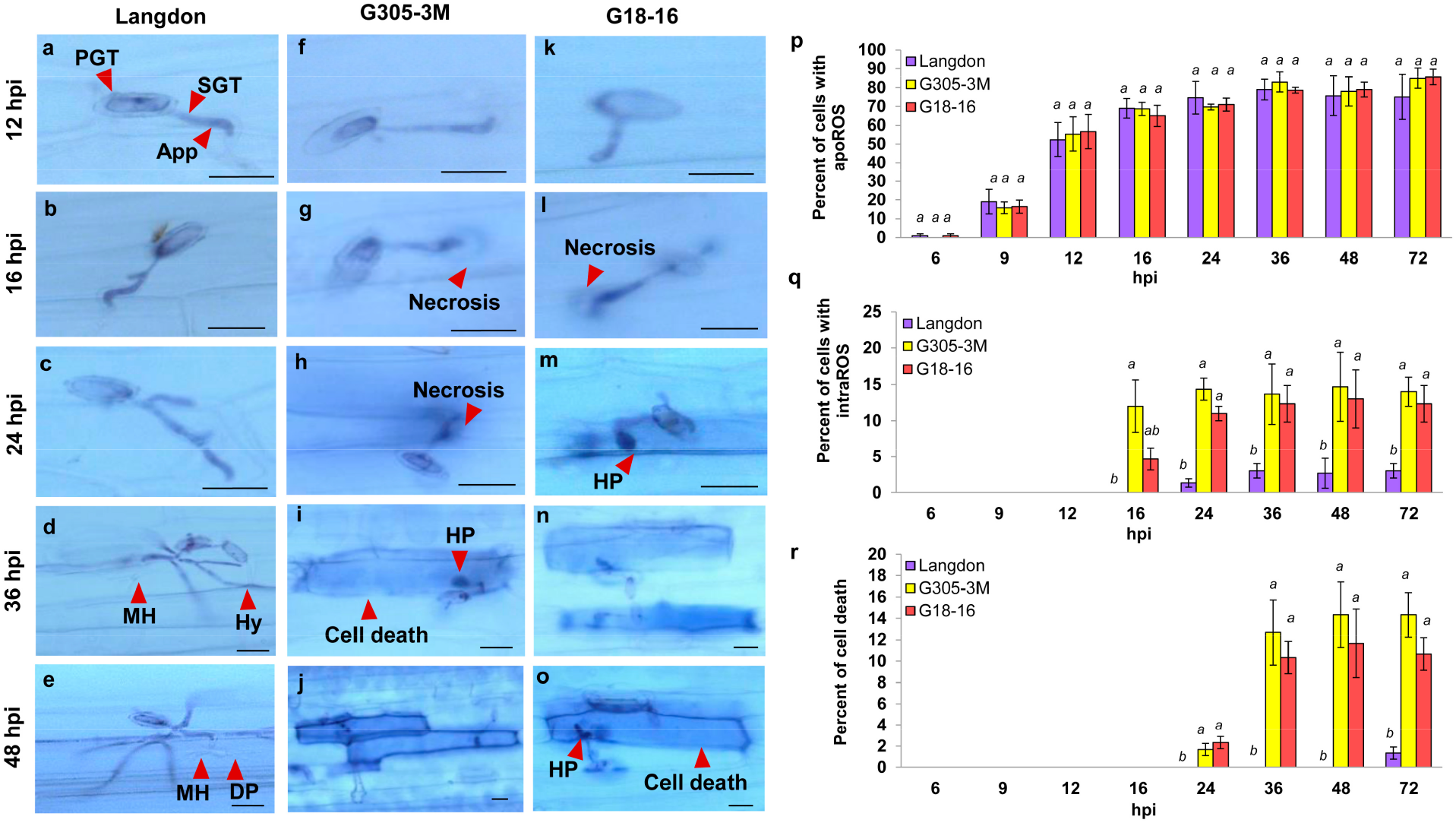
Micrographs of cell death (**a-o**) and quantitative analysis (**p-r**) of the proportion of infected cells with ROS and cell death responses during *Bgt* development in Langdon, G305-3M, and G18-16. App: appressorium; DP, digitate processes; MH, mature haustorium; HP, haustorial primordium; Hy, hyphae; PGT, primary germ tube; SGT, secondary germ tube. Scale bars = 20 μm. (**p**) Percent cells with apoROS of *Bgt*-inoculated cells. (**q**) Percent cells with intraROS of *Bgt*-inoculated cells. (**r**) Percent of cell death of the *Bgt*-inoculated cells. Different letters denote significant differences (p ≤ 0.05) of the mean values by one-way ANOVA analysis to differentiate the accessions at each time piont.

Quantitative assessment of ROS accumulation revealed that at 9 hpi ∼20% of the *Bgt*-infected cells showed apoROS accumulation, then increased to 80% at 36-72 hpi (Fig. 4p). ANOVA analysis showed no significant difference (p≤0.05) between the three genotypes (Langdon, G305-3M and G18-16) at all time points tested (6-72 hpi). In contrast, the intraROS accumulation was detected mainly in the resistant lines (Fig. 4q). IntraROS accumulation began mostly at 16 hpi in G305-3M (∼12%) and G18-16 (∼5%), then it increased to about 11-15% of *Bgt*-infected cells from 24 to 72 hpi duration (Fig. 4q). While in the susceptible Langdon, only 1-4% of the infected cells were detected with intraROS accumulation at 24-72 hpi (Fig. 4q). The cell death response started at 24 hpi in ∼2% of *Bgt*-infected cells and increased to 10-15% after 36 hpi (Fig. 4r), but only in the two resistant WEW lines. The differences in the proportion of infected epidermis cells with intraROS accumulation and cell death response between the resistant and susceptible accessions were highly significant at p≤0.0001 (Fig. 4q, r). In order to validate that the resistance responses were conferred by the *TdPm60* and *Pm69* resistance genes, we have introgressed them into hexaploid common wheat Ruta to obtain BC_4_F_2_ near-isogenic lines (NILs) that harbor these *Pm* resistant genes. These NILs have shown ROS accumulation and cell death resistance responses similar to the *TdPm60* and *Pm69* donor WEW lines, while in Ruta, no intraROS and no cell death were observed (Fig. S2). These results indicate that intraROS-associated HR plays an important role in the *Pm69* and *TdPm60* mediated resistance.

### ROS and cell death in various *Pm* differential lines and among diverse wheat species

To check whether intraROS-associated HR is a common response in resistance to *Bgt*, we characterized these two events in diverse wheat lines that carry various *Pm* genes, and compared them with the response of *Pm69* and *TdPm60* donor lines and NILs (Table 1). We also included in the analysis of six highly susceptible (IT=4) wheat accessions (WEW Zavitan, durum wheat Svevo and Kronos, and bread wheat Morocco Chinese Spring and Ruta). The susceptible lines showed a very low proportion of ROS-mediated cell death response in 0.7-2.3% of the *Bgt-*infected cells, especially in Zavitan and Morocco, while the *Pm* differential lines showed different types/levels of ROS and cell death responses (Table 1, Figs. S3 and S4). *Pm17, Pm41*, and *MlIW72* lines, which are known to carry NLRs, showed strong intraROS and cell death responses, with IT=0-1, that restricted the development of mature *Bgt* haustoria (Fig. S4). The quantitative resistance responses of *MlIW72* were very similar to *TdPm60* (Table 1). *Pm13* and *Pm29* lines induced strong intraROS and cell death, sometimes of 2-3 neighboring cells (Fig. S4). However, while the IT of the *Pm13* line was 1 with only 1.3% developed *Bgt* colonies, the IT of *Pm29* line was 3 (partial susceptibility) with 18.7% developed *Bgt* colonies. For the *pm42* recessive gene line, the ROS and cell death were relatively very low, yet none of the *Bgt* spores developed into a colony, resulting in IT=0. *Pm24* (Wheat Tandem Kinase 3, WTK3) and *Pm32* lines showed partial resistance responses (IT=1-2) with intraROS in 1.6% and 3.3% of *Bgt*-infected cells, respectively. However, the *Pm24* (IT=1) line tended to induce more apoROS accumulation around the penetration peg, with a relatively low percentage of intraROS. *Pm3F* (NLR) line showed very little intraROS and cell death (1-2%) and 52-67.33% of spores that developed *Bgt* colonies, resulting in IT=3-4, suggesting that *Bgt* #70 is virulent on *Pm3F* (Fig. S4). The WEW TD104088 was highly resistant (IT=0) and showed strong ROS and cell death response similar to G305-3M with 15% and 12.3% of *Bgt-*infected cells that showed intraROS accumulation and cell death, respectively (Table 1, Fig. S3). Altogether, intraROS (%) in the *Bgt-*infected cells was positively (r=0.82) and significantly (p≤0.0001) correlated with the cell death response (%) among the different wheat lines (Fig. S5).

An attempt to use H_2_DCFDA for detecting ROS accumulation instead of DAB failed since *Bgt* hyphae also showed fluorescence after the staining with H_2_DCFDA, therefore prohibiting clear detection of apoROS around the penetration peg (Fig. S6).

### Different epidermal cells showed similar ROS and cell death responses

The *Bgt*-mediated ROS and cell death responses were observed all kinds of epidermal cells, including the stomatal guard cells (Fig. S7e, g, k, m), trichomes (Fig. S7i), sister cells (Fig. S7d, f, n, j) and elongated cells (Fig. S7a, b, h, o). These observations suggest that all of these wheat epidermal cells participate in the immune responses activated against *Bgt* which involve apo- and intraROS accumulation and cell death responses.

### *Pathogenesis-related* gene expression patterns in resistant and susceptible wheat

The temporal *PR* gene expression patterns were studied at different time points during powdery mildew infection in G305-3M and Langdon (Fig. S8). The most common patterns were obtained for *PR1* (antifungal), *PR5* (thaumatin-like protein), *PR10* (RNase) and *NPR1* (salicylic acid pathway) genes that showed a peak of expression at 36 hpi in G305-3M, with higher expression levels than in Langdon. The *PR9* (peroxidase) showed higher expression at 48 and 72 hpi in G305-3M than in Langdon, while Langdon showed a peak at 36 hpi. *PR4* (chitinase) and *TaHIR1* (hypersensitive-induced reaction gene) showed a peak of expression at 36 hpi, but the expression levels in Langdon were higher than in G305-3M (Fig. S8). The expressions of *oxacate* gene showed two peaks at 3 and 24 hpi in Langdon, and 9 and 36 hpi in G305-3M, but showing higher expression level in Langdon. Interestingly, *PR14* (lipid-transfer protein) showed a higher expression level in G305-3M before 24 hpi, but lower expression level after 24 hpi relative to Langdon. The expression of *RBOHD* was very low relative to the *PR* genes and the differences between the resistant and susceptible accessions were very small in most of the tested time points (Fig. S8).

### Can ROS accumulation induced by an avirulent *Bgt* isolate provide resistance against a virulent pathogen isolate?

To answer this question, we inoculated wheat cultivar Morocco with a mixture of an avirulent isolate *Bgt* #SH that induces ROS accumulation and a virulent isolate *Bgt*#70 that can cause disease (Fig. 5a-b). However, although ROS was successfully induced by the avirulent isolate, it did not prevent the development of *Bgt* colonies (Fig. 5c). In a second trial, we inoculated Morocco first with the avirulent *Bgt* #SH, and after 16 hpi (intraROS burst time) with the virulent isolate *Bgt* #70. However, Morocco still showed a highly susceptible phenotype (IT=4), as in single inoculation with *Bgt* #70 (Fig. 5d), suggesting the intraROS induced by the avirulent *Bgt* isolate #SH did not induce effective resistance against a virulent *Bgt* isolate.

**Fig. 5.**
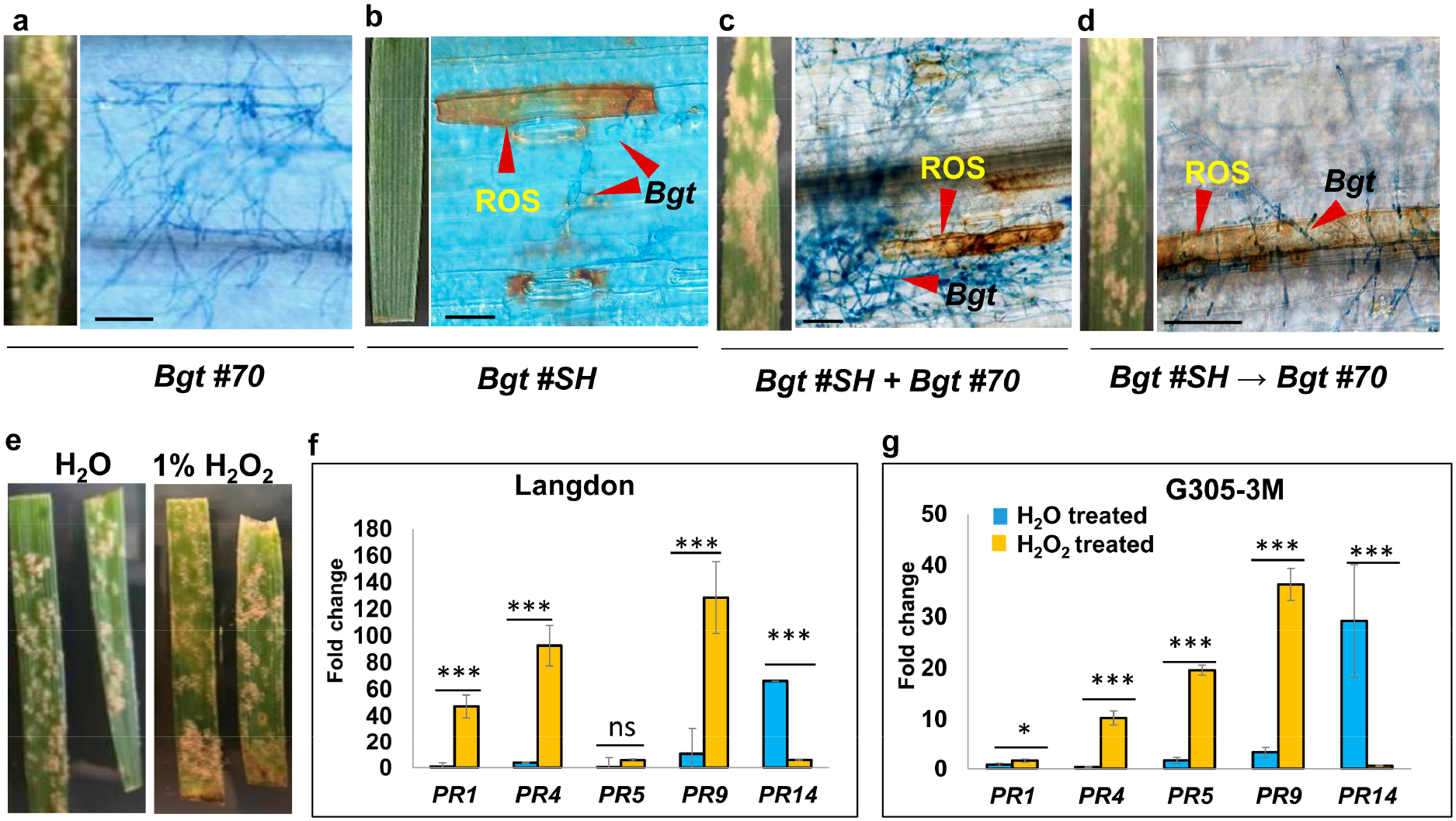
ROS accumulation is not a strong SAR signal. **(a)** The phenotypes (at 7 hpi) and representative micrograph (at 48 hpi) of Morocco to inoculation of single *Bgt* #70 (IT=4), **(b)** single *Bgt* #SH (IT=0), **(c)** both isolates (*Bgt* #70 + *Bgt* #SH) (IT=4), and **(d)** first *Bgt* #SH then *Bgt* #70 after 16 hours (*Bgt* #SH → *Bgt* #70). Scale bars = 100 μm. **(e)** The phenotypes of Langdon to *Bgt* #70 (both IT = 4) with pretreatments of H_2_O or 1% H_2_O_2_ before *Bgt* infection. The expression levels of *PR* gene in water (H_2_O) and H_2_O_2_-treated (24 hours after treatment) leaves of Langdon (**f**) and G305-3M (**g**). Asterisks indicate the level of significance by *t-*test, p ≤ 0.05 (*), p ≤ 0.01 (**), p ≤ 0.001 (***), non-significant (*ns*) to differentiate the *PR* genes in response to treatments (H_2_O and H_2_O_2_).

Moreover, 1% H_2_O_2_ treatment was not able to induce resistance to *Bgt* #70 in Langdon (Fig. 5e), although it significantly upregulated the expressions of some important *PR* genes (*PR1, PR4, PR5* and *PR9*) (Fig. 5f-g). Altogether, these results were suggesting that ROS could not act as a strong systemic signal for inducing high resistance to *Bgt* in wheat.

## DISCUSSION

In this study, the contribution of ROS (intraROS and apoROS) and localized cell death to the immune responses were investigated quantitatively in the wheat-powdery mildew pathosystem, in the presence of various *Pm* genes. In most cases, hypersensitive cell death response followed intraROS accumulation as part of the resistance mechanism activated by NLR genes, while in some unconventional *Pm* genes, different immune responses were observed. Furthermore, we demonstrated that the ROS accumulation activated by an avirulent isolate did not induce resistance against a virulent isolate.

### ROS accumulation in response to powdery mildew infection

ROS accumulation is necessary for the activation of plant immunity and the regulation of resistance mechanisms (Hu et al. 2021; Mittler 2017). In the current study, we were able to quantitatively differentiate between two types of spatial ROS accumulation in response to *Bgt* infection, the apoROS and the intraROS, which are probably activated by PTI and ETI, respectively (Halliwell 2006; Hückelhoven 2007; Marti et al. 2021). The apoROS is known to be secreted by the membrane-bound RBOHD activated by receptor-like kinases in response to the detection of chitin (a component of the fungal cell wall) or plant cell wall degradation products upon the pathogen infection (Lee et al. 2020). In the current study, we detected apoROS around the PGTs in about 70-80% of the infected cells, both in the resistant and the susceptible genotypes, already at 16 hpi, with no significant difference between them ((p≤0.05, Fig. 4p). Yamaoka et al. (2007) have shown that the primary germ tubes of *Blumeria graminis* are involved in the suppression of resistance induction of host plant cells. Therefore, it seems that apoROS by itself is not sufficient to prevent disease development caused by *Bgt*. Although some ROS could be observed around the haustorium primordia (HP) of *Bgt* in the compatible interactions during the early infection stages (16-72 hpi) (Fig. 2d, e, g, k, l), no intraROS accumulation was found at later stages around the mature haustoria (MH) (Fig. 2e, i). These results may indicate that MH are able to suppress intraROS accumulation probably by secretion of effectors into the host cytoplasm (Jwa and Hwang 2017; Liu et al. 2021). Previously, it was shown that apoROS might enter the cytoplasm through endocytosis or membrane-bound aquaporin channels (Mittler et al. 2022; Rodrigues et al. 2017). These findings may explain our results showing that some intraROS may appear when extensive apoROS accumulation occurs in the compatible interaction (Fig. 2e, n and p). A high proportion of *Bgt*-infected cells with intraROS (∼15%) was observed only in the resistant genotypes (Fig. 3d-g, 3n-q), but not in the susceptible genotypes. These results probably represent intraROS accumulation induced by ETI as a result of the recognition of *Bgt* effectors by host specific genes (Dalio et al. 2021). The main producers of intraROS are probably the chloroplast, mitochondria, and/or peroxisome as a consequence of the imbalance and disruption of metabolic pathways during plant-pathogen interactions (Camejo et al. 2016; Littlejohn et al. 2021; Mittler et al. 2022; Su et al. 2018). An example of such immune response was demonstrated for the Wheat Kinase Start1 (*WKS1*) stripe rust resistance gene (*Yr36*) which was shown to increase chloroplast H_2_O_2_ accumulation by phosphorylation of the thylakoid-associated ascorbate peroxidase causing accumulation of ROS and cell death (Gou et al. 2015).

### The resistance cell death response

HR mediated cell death has been known to block (hemi)biotrophic pathogen colonization through the signaling pathways triggered by host plant’s NLRs mediated recognition of pathogen effectors (Pitsili et al. 2020). The PRRs signaling of PTI may be monitored by NLRs, with PRR signaling disturbance leading to hypersensitive cell death response (Pitsili et al. 2020). PTI is required for full induction of ETI and in turn that ETI induces and stabilizes key PTI signaling components (Bjornson and Zipfel 2021). In our study, the cell death response was mainly recorded in the resistant genotypes following intraROS accumulation at a delay of ∼20 hours. However, in the susceptible wheat genotypes, the apoROS triggered by PTI did not lead to cell death responses (e.g. Fig. 2 and 4). The spatial distribution of intraROS and cell death responses were observed in all types of epidermal cells (Fig. S7), including stomatal guard cells, trichomes, sister cells, and elongated cells. Therefore, our results support previous reports showing that intraROS accumulation plays an important role in NLR-mediated cell death (Bjornson and Zipfel 2021; Dalio et al. 2021). Interestingly, around 80% of *Bgt*-infected cells in the resistant genotypes showed obvious accumulation of apoROS, while no intraROS accumulation was observed (Table 1). These results may indicate that *Bgt* infection was probably restricted already in the PTI stage., thus supporting previous results indicating the crosstalk between PTI and ETI immune receptors is involved in the plant immune responses (Bjornson and Zipfel 2021). Moreover, recent studies show that pathogen infection can trigger NLR receptors to form a macromolecular porous structure called resistosome participating in the cell death signaling hubs. The cell death is preceded by a perturbation of organelle hemostasis and channel-dependent ROS production, followed by a loss of plasma membrane integrity (Bi et al. 2021; Wang et al. 2019). Therefore, our results are in agreement with previous studies and confirm that intraROS production plays important role in NLR-mediated cell death, also in the *Bgt*-wheat pathosystem.

### Differential *Pm* genes conferred different resistance responses

ROS accumulation and HR cell death immune responses of differential wheat lines that harbor various *Pm* resistance genes were compared in the current study. Based on the host plant’s immune responses, the *Pm* genes harbored by these lines can be classified into three different categories: (A) Typical NLR responses which include *Pm69* (Li et al. 2022), *TdPm60, MlIW72, Pm3F* and *Pm41* with IT=1-3. These different levels of defense responses might be due to the different interactions of the *Bgt* #70 effectors with the specific NLRs (Dalio et al. 2021). The resistance levels provided by these NLRs were strongly associated with intraROS and cell death levels (Table 1, Fig. S5), supporting previous studies showing that the intraROS and cell death play an important role in NLR-mediated resistance (Bjornson and Zipfel 2021; Dalio et al. 2021). (B) The *pm42* is a recessive resistance gene (Hua et al. 2009), which did not induce a strong intraROS burst nor cell death response (Table 1), suggesting a different resistance mechanism compared with the typical NLRs. A resistance response conferred by a recessive gene may indicate the lack of a cell component necessary for the pathogen to proliferate or the lack of a negative regulator of plant immunity pathways (Deslandes et al. 2002). (C) *Pm24* is encoding for a Wheat Tandem Kinase protein (WTK3) that belongs to a newly discovered family of intracellular disease resistance proteins, which also activate cell death responses (Klymiuk et al. 2021). *Pm24* governs only partial resistance (IT=1) to *Bgt* (Table 1, Fig. S4). In accordance with Lu et al. (2020), our results demonstrate that only ∼3.3 % of *Pm24* cells infected with *Bgt* showed intraROS accumulation, while ∼10% of *Bgt*-infected cells presented cell death response, which suggests that a new disease resistance mechanism is involved here, which is probably different from that of NLRs. Klymiuk et al (2021) proposed that tandem kinases are new players in the plant immune system serving as decoys for pathogen effectors and activating PCD.

### ROS-induced *PR* gene expression and is not a strong SAR signal

Several important signals have been reported to be involved in plant defense responses, especially in systemic acquired resistance (SAR) that induces systemic production of antimicrobial proteins known as PR proteins after pathogen infection (Fu and Dong 2013; Li et al. 2020; Phuong et al. 2020). In the current study, *Bgt* infection induced a temporal expression pattern of *PR* genes (*PR1, PR4, PR5, PR9, PR10* and *PR14*) (Fig. S8), suggesting that those *PR* genes are involved in wheat immune responses. We also showed that external ROS application could upregulate *pathogenesis-related* (*PR*) genes, but not induced high resistance for the whole wheat leaf (Fig. 5e). IntraROS induced by an avirulent *Bgt* isolate did not contribute to an induced resistance against a virulent isolate in the same wheat leaf (Fig. 5). This study is paving the way for future studies aiming to dissect the disease resistance mechanism of diverse wheat genetic resources against *Bgt* at the molecular level.

## ACKNOWLEDGEMENTS

Yinghui Li is thankful for the postdoctoral fellowship (2018-21) provided by the Planning and Budgeting Committee (PBC) of the Israel Council for Higher Education (ICHE) for Outstanding Postdoctoral Fellows from China and India. Rajib Roychowdhury is thankful for the combined Post-Doctoral Fellowship during 2020-21 provided by the Graduate School of the University of Haifa, Israel. The authors are thankful to Dr. Tamar Lotan laboratory at the University of Haifa for assistance in microscopy work, and to Dr. Tamar Kis-Papo and Dr. Olga Borzov for technical assistance. This study was supported by the Israel Science Foundation, grant numbers 1366/18, 2289/16 and 2342/18.

## The authorship contribution statement

**TF, YL** and **RR** conceived and designed this research; **YL** and **RR** performed the experiments and data analysis; **YL, RR** and **TF** wrote the manuscript; **LG** and **SJ** assisted in laboratory experiments; **ZW, IS** and **TF** proof-read, reviewed and edited the manuscript and improved it with additional suggestions; **TF** was responsible for coordination and funding acquisition. All the authors approved the final version of the manuscript for submission for publication.

## Supplementary Information

Figures S1-S8 and Table S1. See the attached Supplementary file.

