## Supplemental for "Intracellular reactive oxygen species (intraROS)-aided localized cell death contributing to immune responses against wheat powdery mildew pathogen"

**Figure S1-S8 and Table S1.**

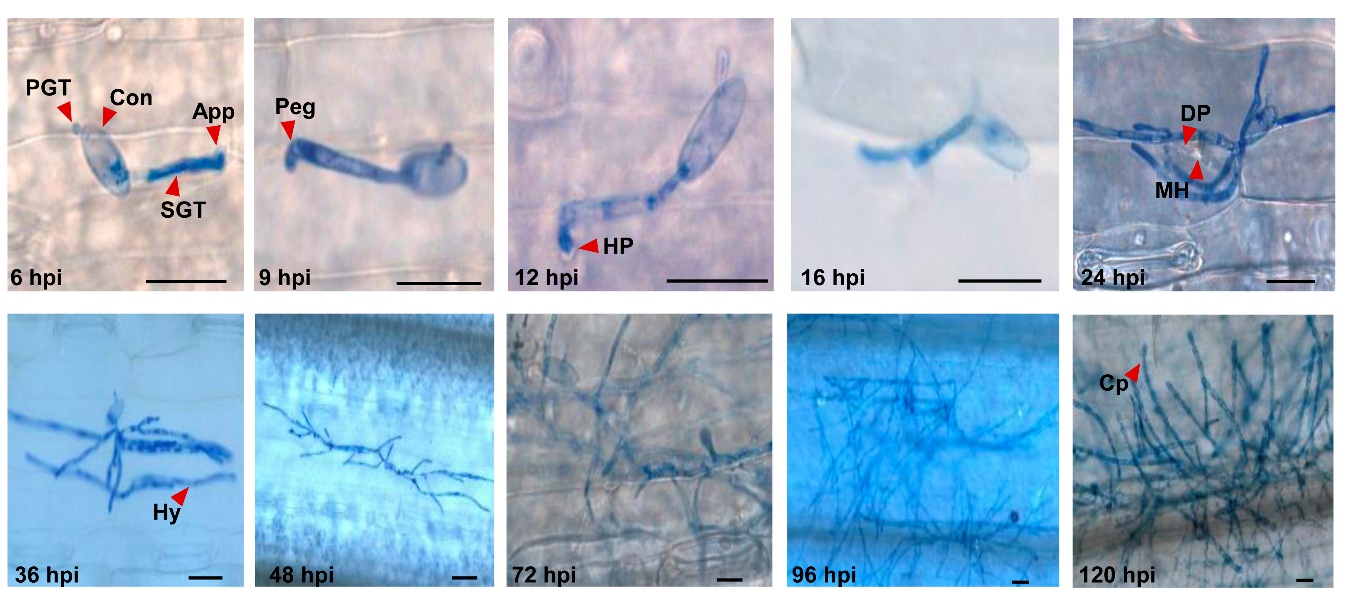

**Fig. S1.** Representative micrograph of *Bgt* #70 development on susceptible Langdon without ROS accumulation. The spores and fungal structures are colored in blue. App, appressorium; Con, conidium; Cp, conidiophore; DP, digitate processes (finger-like projections); MH, mature haustorium; HP, haustorial primordium; Hy, hyphae; Peg, penetration peg; PGT, primary germ tube; SGT, secondary germ tube. Scale bars = 100 μm.

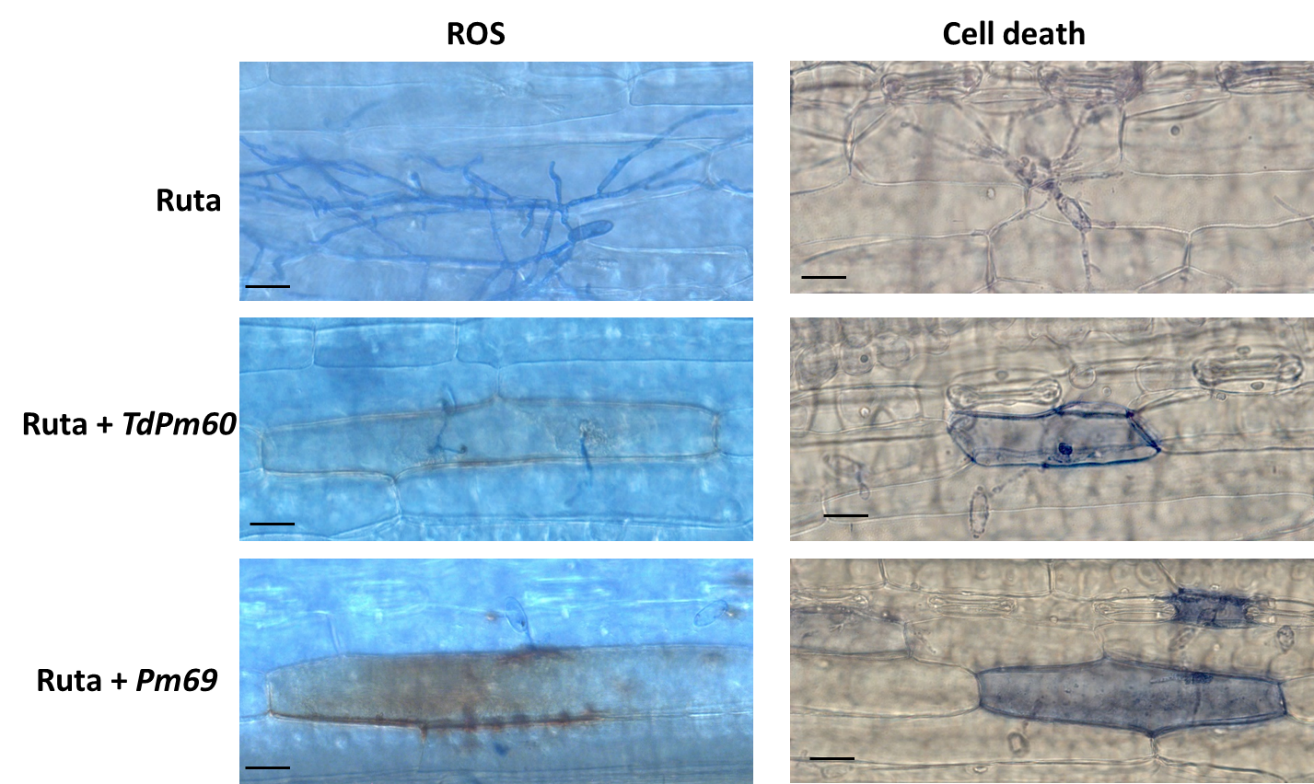

**Fig. S2.** Representative micrograph showing ROS accumulation and cell death for *Bgt*#70 infection in Ruta and its two introgression lines (ILs) harboring powdery mildew resistance genes- *TdPm60* and *Pm69*, individually. Scale bars = 20 μm.

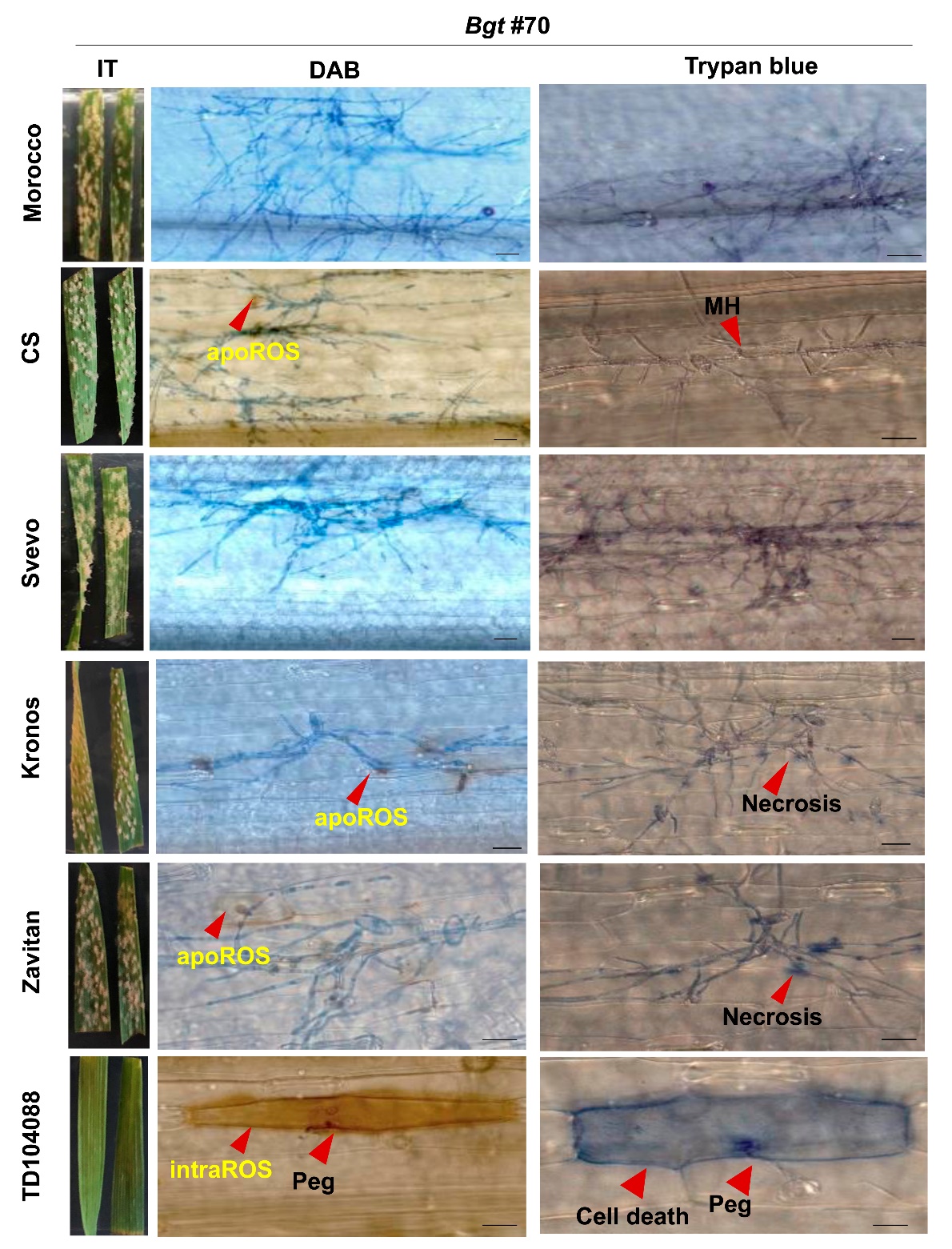
**Fig. S3.** Representative micrograph of ROS accumulation (brown colored cells) and cell death (blue colored cells) in response to *Bgt* #70 in different wheat accessions. Bread wheat accessions: Morocco and Chinese Spring (CS); Durum wheat accessions: Svevo and Kronos; WEW: Zavitan and TD104088. The spores and fungal structures are also colored in blue. Scale bars = 100 μm.

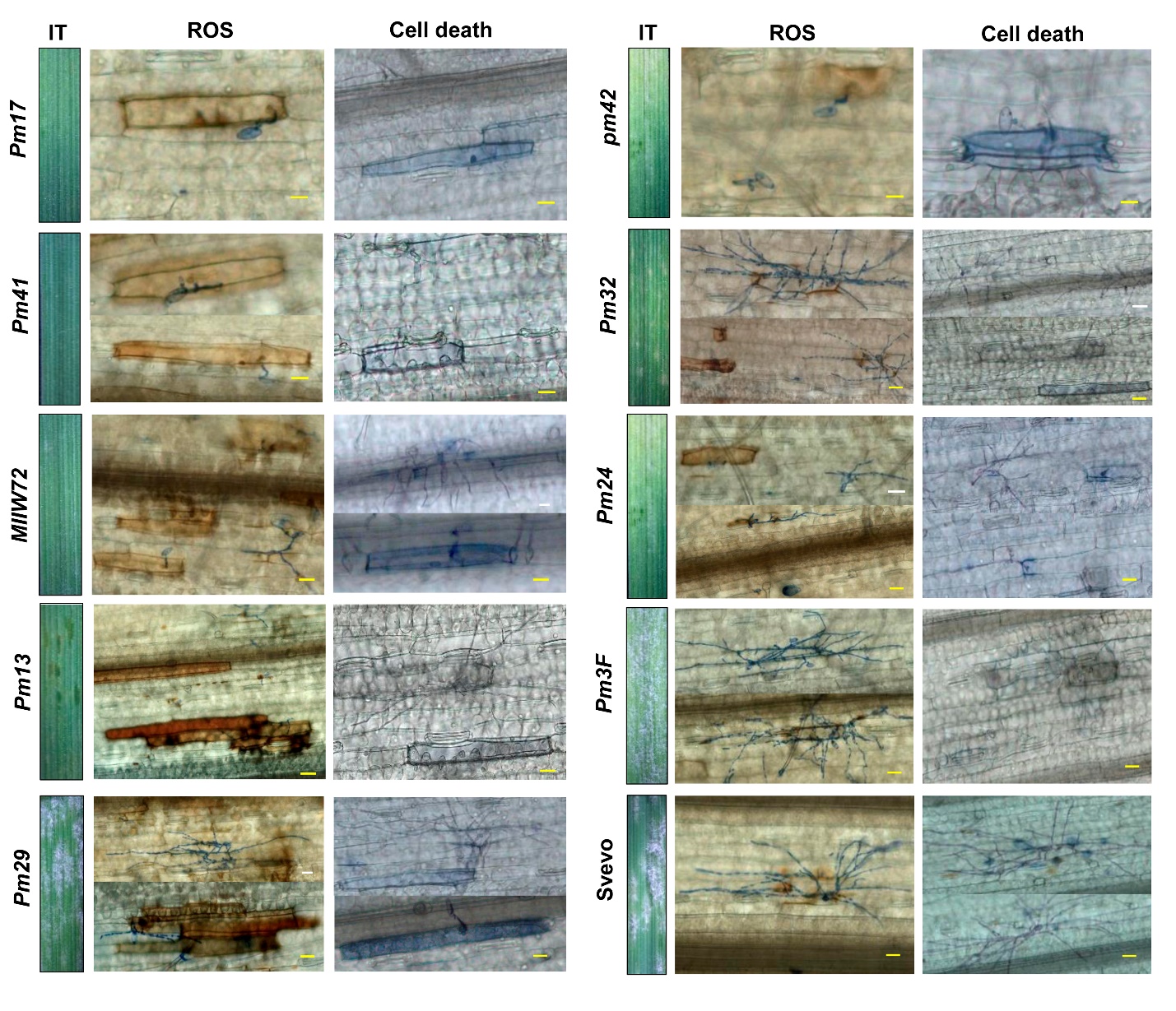

**Fig. S4.** Representative micrograph of macroscopic (phenotype) and microscopic (ROS accumulation and cell death) observations in *Pm* differential wheat lines and durum wheat (Svevo, as susceptible control) in response to *Bgt* #70 at 48 hpi. Scale bars = 20 μm.

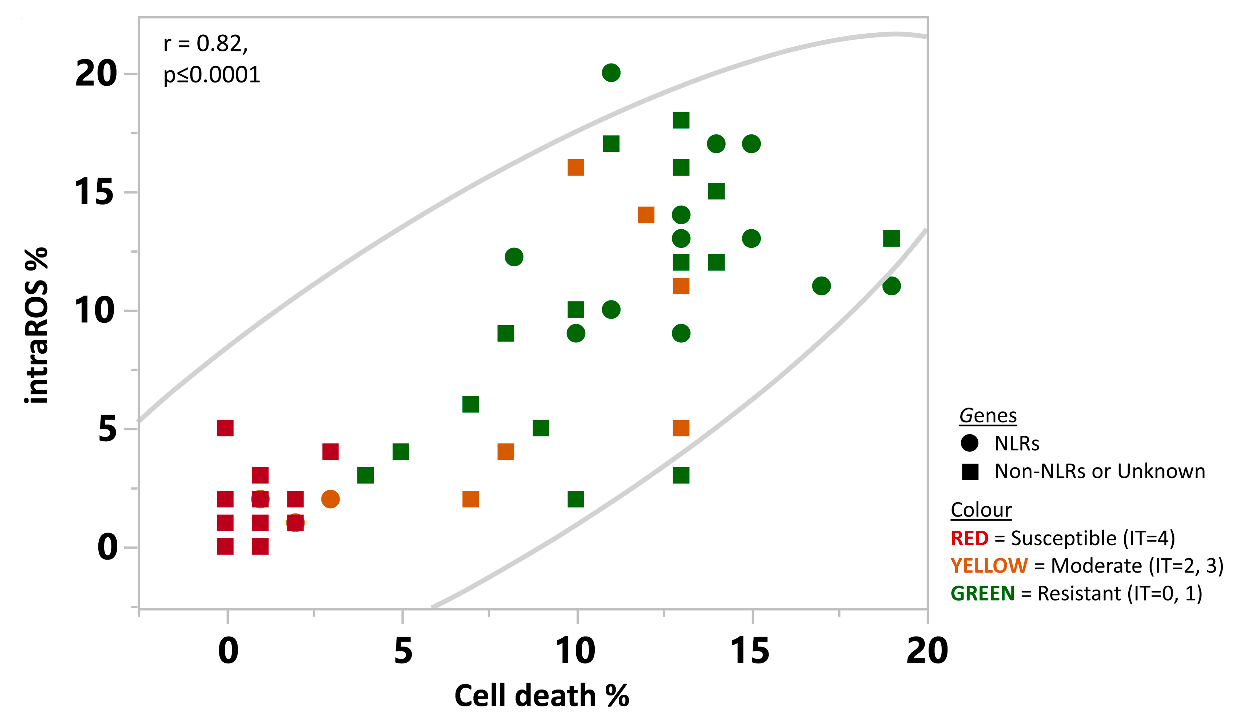

**Fig. S5.** A correlation analysis between intraROS% and cell death% in the *Bgt*-infected cells among different wheat accessions and differential lines**.** Different marker shape represents different *Pm* genes and different colors denote the level of resistance/susceptibility to *Bgt* #70.

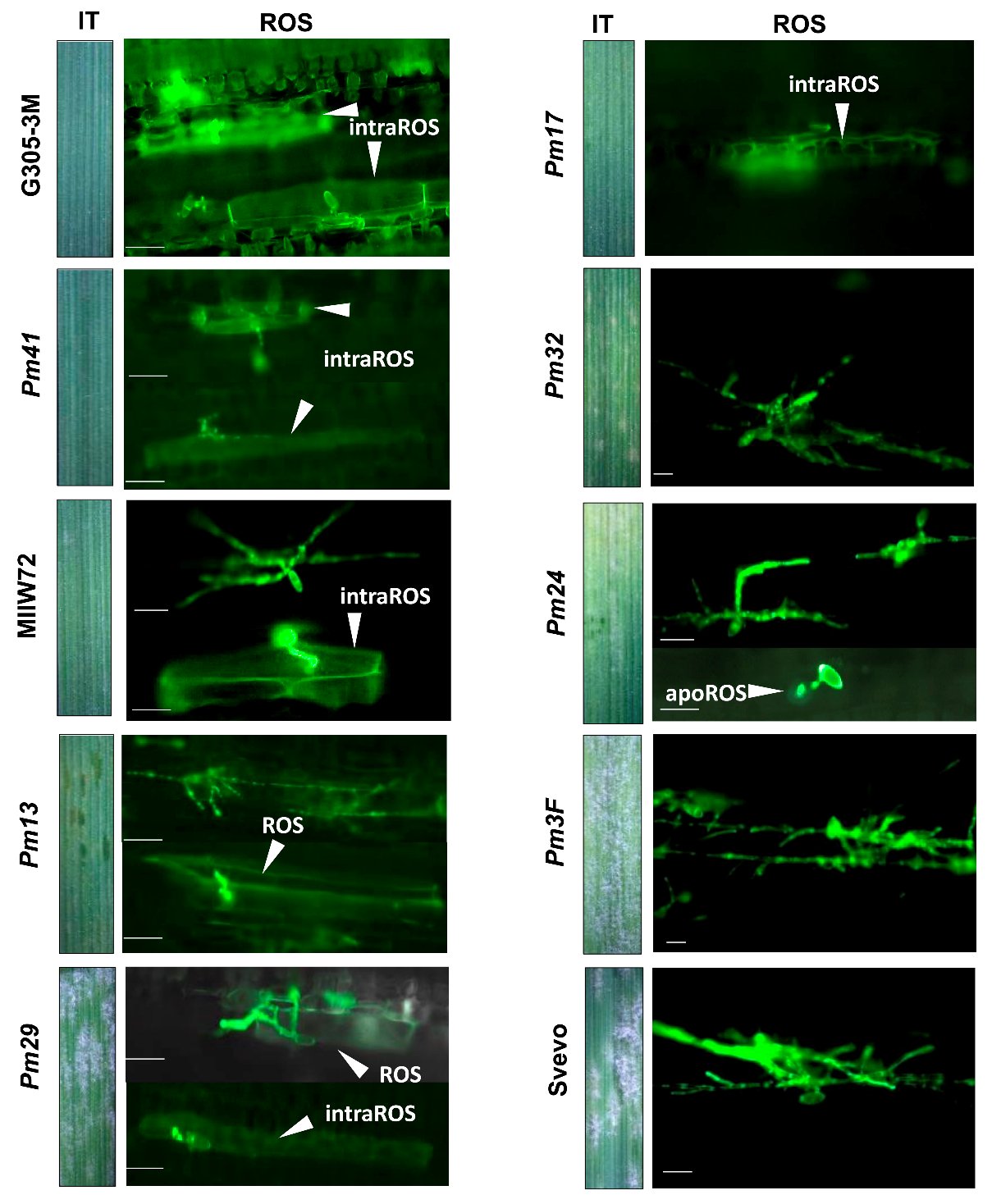
**Fig. S6.** Representative micrograph of macroscopic (phenotyping white *Bgt* colonies) and microscopic (ROS accumulation as green fluorescence) observations in *Pm* differential wheat lines, WEW (G305-3M) and durum wheat (Svevo) in response to *Bgt* #70. Cellular ROS and fungal structures were observed under a confocal microscope using 2′,7′-dichlorofluorescein diacetate (H_2_DCFDA), and cell death was observed under a light microscope. Scale bar = 100 μm.

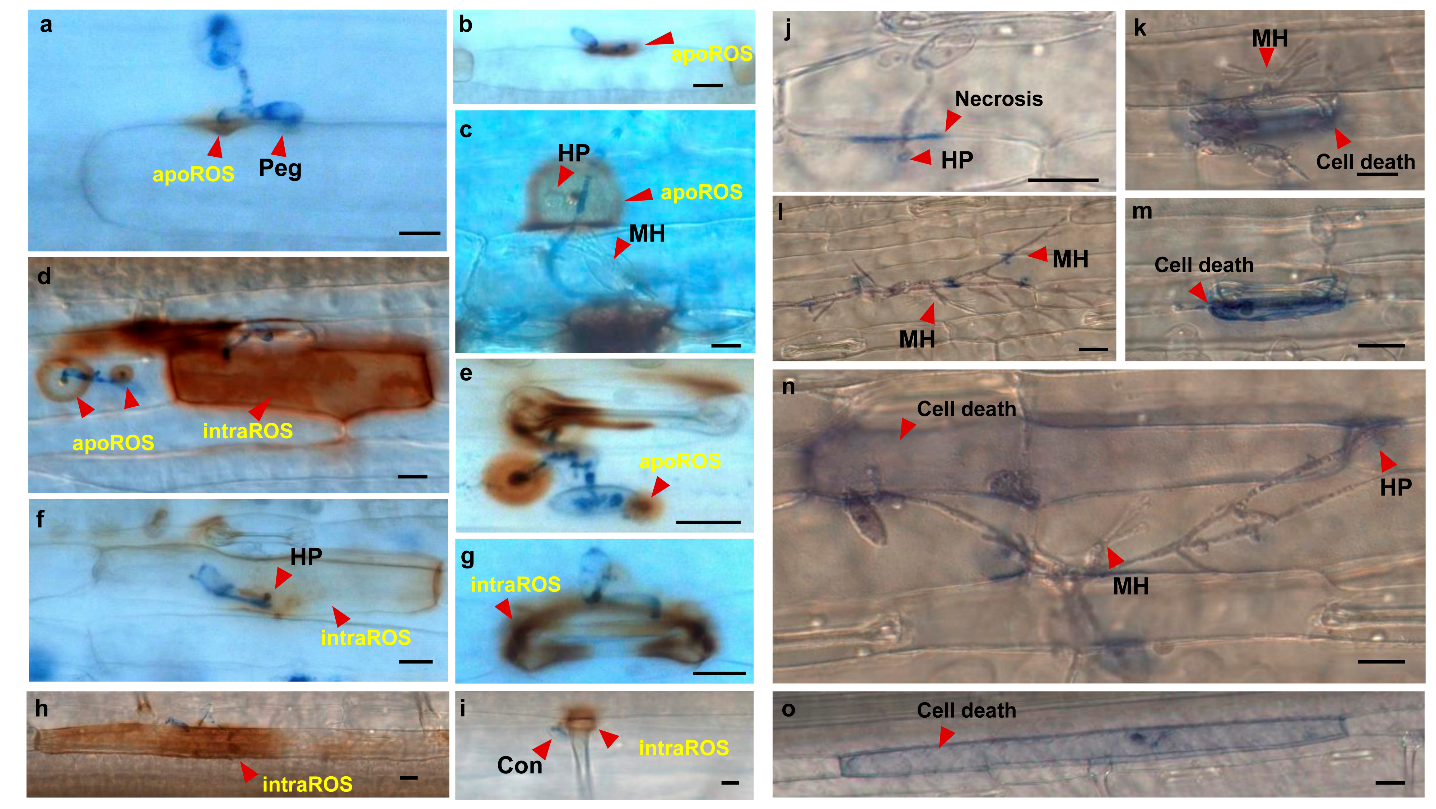

**Fig. S7.** Representative micrograph of ROS (reddish-brown coloration) and cell death (blue coloration) response in different kind of wheat cells. The spores and fungal structures are colored in blue. Stomatal guard cells (**e, g, k, m**), trichomes (**i**), sister cells (**d, f, n, j**), elongated cells (**a, b, h, o**). (**c, h, i**) TD104088 with *Bgt* #SH at 48 hpi. (**a, b, e**) G305-3M with *Bgt* #70 at 48 hpi. (**d**) G305-3M with *Bgt* #70 at 16 hpi. (**f**) G18-16 with *Bgt* #70 at 36 hpi. (**g**) G18-16 with *Bgt* #70 at 48 hpi; (**j**) Ruta with *Bgt* at 16 hpi. Scale bars = 20 μm.

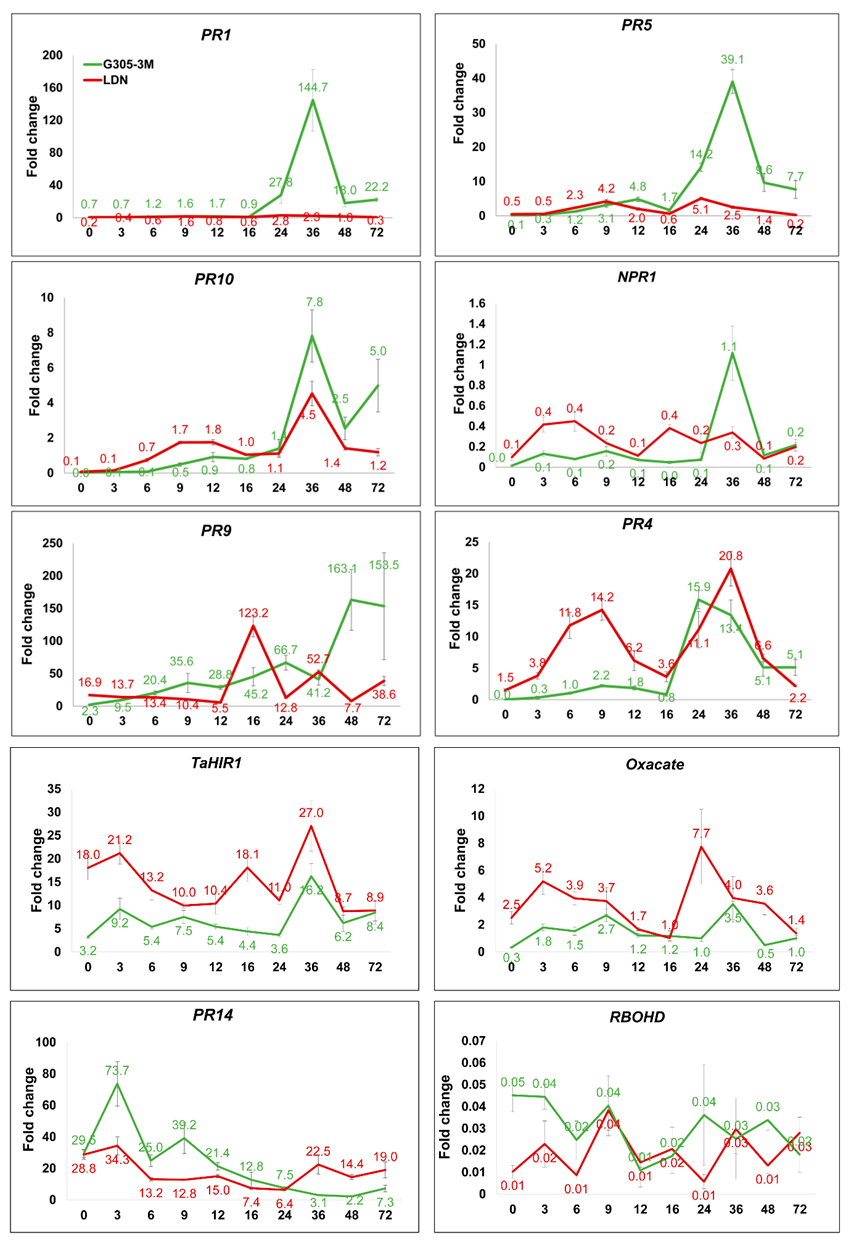

**Fig. S8.** Expression patterns of different *PR* genes at different points in infection time for resistant G305-3M and susceptible Langdon. X-axis is the *Bgt* #70 inoculation time: 0-72 hpi (hours post infection). Y-axis is a relative expression of the *PR* gene (Target/Actin gene).

TABLE S1. A list of *PR* and other genes involved in plant immune response used for q-PCR, together with DNA primer sequences, size of expected products, and description of the genes.

| **Name of the gene** | **Primer sequences (5´-3´) for qPCR** | **PCR product Size (bp)** | **Tm (ºC)*^1^*** | **Description of genes** |
| --- | --- | --- | --- | --- |
| *Actin* | F: ACCTTCAGTTGCCCAGCAAT  R: CAGAGTCGAGCACAATACCAGTTG | 91 | 62  63 | TRIDC1BG046410.5 actin 7 |
| *PR1* | F: CTGGAGCACGAAGCTGCAG  R: CGAGTGCTGGAGCTTGCAGT | 76 | 64  66 | TRIDC7AG019820.1 Cysteine-rich venom protein (antifungal) |
| *PR4* | F: CGAGGATCGTGGACCAGTG  R: GTCGACGAACTGGTAGTTGACG | 128 | 63  63 | TRIDC3BG084520.2 Wound-induced protein (chitinase) |
| *PR5* | F: CAGACTACTACGACATCTCGG  R: ACTGAACCTCCTGTTACCG | 154 | 60  60 | TRIDC5AG002750.3 Thaumatin-like protein |
| *PR9* | F: GACTGTTCCACTTGGGAGAC  R: CGACCATGTCGACCGTGTTA | 140 | 60  62 | RIDC2BG015920.2 Peroxidase superfamily protein |
| *PR10* | F: GGCCGTTCGATAAGTTCCA  R: CTTGAAGGGGTGGAAGAGGA | 128 | 60  61 | TRIDC1BG028040.1 Ribonuclease T2 family protein |
| *PR14* | F: GAGCCCCTGCATCTCCTATG  R: GCTGGCTAGACTCCTAACGC | 82 | 63  63 | TRIDC0UG009250.1 Non-specific lipid-transfer protein 4.1 |
| *Oxalate oxidase* | F: GCTCTACTCCAGGGTGGTGC  R: TGGCTGTTGAAGGAGACGAC | 117 | 66  62 | TRIDC4BG005640.1 germin-like protein 4 |
| *NPR1* | F: AGCCCTCTCCAAAACAGTCG  R: GATGCATCTCTTCCGAGCGA | 114 | 62  62 | TRIDC3AG012960.1 regulatory protein (NPR1) |
| *TaHIR1* | F: ACAGGCTTAGCAACACCAGG | 162 | 63  62 | TRIDC5AG070770.3 hypersensitive-induced response protein 1 |
|  | R: CGTACCCGTAGGCAGACA |  |  |  |
| *RBOHD* | F: GCTCAAGCACATCTTCGTC  R: GTCGTAGTTCCTGAAATCCTG | 174 | 60  60 | TRIDC6BG031500.2 respiratory burst oxidase homologue D |
| ***^1^*** Annealing temperature is ranged around in general 59-60 ºC for all used *PR* genes for qPCR. | | | | |
